# A heterologous in-cell assay for investigating intermicrovillar adhesion complex interactions and function

**DOI:** 10.1101/2020.09.06.285171

**Authors:** Meredith L. Weck, Scott W. Crawley, Matthew J. Tyska

**Affiliations:** Department of Cell and Developmental Biology, Vanderbilt University, Nashville, TN 37232; Department of Biology, The University of Toledo, Toledo, Ohio

**Author notes:** **To whom correspondence should be addressed:** Matthew J. Tyska, Ph.D., Department of Cell and Developmental Biology, Vanderbilt University School of Medicine, T-2212 Medical Center North, 465 21st Avenue South, Nashville, TN 37240-7935.

**Keywords:** filopodia, microvilli, brush border, enterocyte, epithelia, actin, bundle, cytoskeleton, myosin, protocadherin, MYO10, MYO7B, CDHR2, CDHR5, USH1C, ANKS4B

## Abstract

Solute transporting epithelial cells build arrays of microvilli on their apical surface to increase membrane scaffolding capacity and enhance function potential. In epithelial tissues such as the kidney and gut, microvilli are length-matched and assembled into tightly packed ‘brush borders’, which are organized by ∼50 nm thread-like links that form between the distal tips of adjacent protrusions. Composed of protocadherins CDHR2 and CDHR5, adhesion links are stabilized at the tips by a cytoplasmic tripartite module containing the scaffolds USH1C and ANKS4B, and the actin-based motor, MYO7B. As several questions about the formation and function of this ‘intermicrovillar adhesion complex’ remain open, we devised a system that allows one to study individual binary interactions between specific complex components and MYO7B. Our approach employs a chimeric myosin consisting of the motor domain of MYO10 fused to the cargo-binding tail domain of MYO7B. When expressed in HeLa cells, which do not normally produce adhesion complex proteins, this motor exhibited robust trafficking to the tips of filopodia and was also able to transport individual components to these sites. Unexpectedly, the MYO10/MYO7B chimera was able to deliver CDHR2 and CDHR5 to distal tips in the absence USH1C or ANKS4B. Cells engineered to localize high levels of CDHR2 at filopodial tips acquired inter-filopodial adhesion and exhibited a striking dynamic length matching activity that aligned distal tips over time. These observations reveal a robust adhesion-dependent mechanism for matching the lengths of adjacent surface protrusions, and may offer insight on how epithelial cells minimize microvillar length variability.

## INTRODUCTION

All cells use plasma membrane protrusions to physically and biochemically interact with the external environment. Finger-like protrusions such as microvilli, stereocilia, and filopodia extend from the surface of diverse cell types and serve a variety of functions, ranging from solute uptake to mechanosensation (1). Individual protrusions are supported by parallel bundles of actin filaments, with their barbed ends oriented away from the cell body toward the distal tip. Microvilli in particular play prominent roles in transporting epithelial tissues, including those that line the intestinal tract and kidney tubules (2,3). For example, a single enterocyte, the nutrient absorbing cell of the gut, assembles on its apical surface thousands of microvilli in an array collectively known as the brush border (BB). The BB vastly increases surface area and in turn, absorptive capacity (4). In this context, microvilli exhibit near perfect packing organization and very low length variability (5); the ∼2500 microvilli that extend from the apical surface of a single enterocyte exhibit less than 5% length variability (6). Such length matching gives rise apical surface flatness and may be important for protecting these cells from mechanical damage induced by flow or peristaltic compression (7). Although molecular mechanisms of microvillar organization have begun to emerge in recent years (7–13), how microvillar length is controlled or matched to the dimensions of adjacent protrusions remains unclear.

Recent cell biological studies have revealed that the packing organization of epithelial microvilli is constrained by extracellular adhesion links. Ultrastructural imaging of native intestinal tissues showed that the tips of individual microvilli are connected to their neighbors by 12-15 radially arranged links ∼50 nm in length, which are at least partially composed of CDHR5 and CDHR2 ectodomains interacting in *trans* to form strong heterophilic complexes (11). CDHR2 ectodomains can also interact in *trans* to form weaker homophilic complexes, although the biological significance of weak homophilic vs. strong heterophilic adhesion remains unclear (11). CDHR2 and CDHR5 enrichment at microvillar tips is promoted by interactions with cytoplasmic binding partners, including the two scaffolding proteins, ankyrin repeat and sterile a motif domain containing 4B (ANKS4B) and harmonin/Usher syndrome 1C (USH1C), as well as the actin-based motor, myosin-7b (MYO7B) (9–11,13). Most recently, CALML4 was identified as a small calmodulin-like molecule that likely serves as an activity-supporting light chain for MYO7B (14). Together with the microvillar protocadherins, we refer to the complex formed by all of these proteins as the ‘intermicrovillar adhesion complex’ (IMAC, Fig. 1A). A similar ‘tip-link’ complex, consisting of USH1C, SANS (related to ANKS4B), MYO7A (related to MYO7B), PCDH15 (related to CDHR5) and CDHR23 (related to CDHR2), is found at the distal ends of stereocilia on the surface of hair cells in the inner ear (12,15–26). Here, the tip-link complex plays a critical role in mechanosensation, and loss-of-function mutations in any tip-link molecule lead to Usher syndrome, which is characterized by progressive deaf-blindness (27,28). USH1C is the only factor expressed in both inner ear and intestinal adhesion complexes, and mutations in this gene lead to functional and morphological defects in both tissues, in humans and mice (29,30). The appearance of similar distal tip-linking adhesion complexes in distinct tissues with remarkably different physiological functions suggests that these multiprotein complexes fulfill a fundamental role in epithelial apical morphogenesis (31).

**Figure 1:**
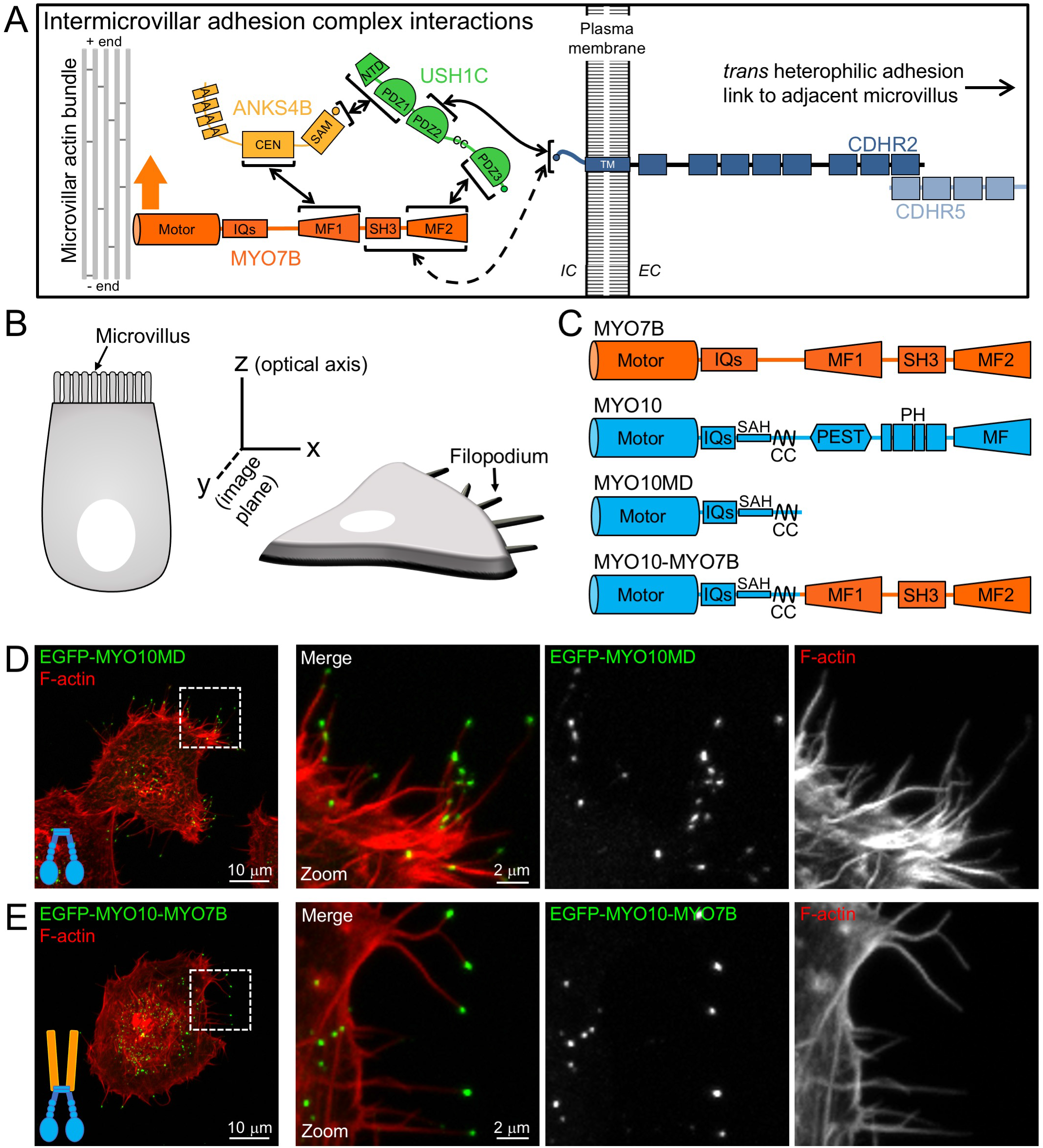
Strategy for studying IMAC component interactions in a heterologous system using a chimeric myosin. (A) Cartoon highlighting the numerous interactions between individual cytoplasmic elements of the IMAC. Solid arrows represent well characterized binary interactions that have been defined at the atomic level; dashed arrows represent interactions identified with pulldowns but lacking structural information. (B) Cartoon illustrating the orientation of finger-like protrusions extending from the surface of a polarized epithelial cell (*left*) vs. a non-polarized HeLa cell (*right*); HeLa filopodia extend parallel to the image plane rather than the optical axis. (C) Domain organization of MYO7B, MYO10, and the truncated (MYO10MD) and chimeric (MYO10-MYO7B) myosin constructs used in these studies. (D) Confocal image of HeLa cells expressing EGFP-MYO10MD (green), which lacks the WT MYO10 tail domain but retains filopodial tip targeting potential; cells are counterstained with phalloidin to label F-actin (red). The white dashed outlined box in the overview panel at left indicates the region shown in the merge and single channel zoom panels to the right. (E) Confocal image of HeLa cells expressing EGFP-MYO10-MYO7B (green) and counterstained with phalloidin to label F-actin (red). Dimensions of scale bars are indicated in figure panels.

Previous studies established that each IMAC component is required for normal BB formation; results from KD of MYO7B, CALML4, ANKS4B, CDHR2 or CDHR5 in cultured epithelial cells (9–11,14), and KO of CDHR2 or USH1C in mice (7,11) collectively show that removal of a single complex component reduces the levels of all other components and leads to significant defects in microvillar organization and BB morphology. Moreover, biochemical and structural studies have defined the binding properties and atomic scale details of most of the binary interactions between IMAC proteins (8,12,13). One theme that emerges is multivalence; that is, all IMAC components interact with at least two and in some cases three other components (Fig. 1A). MYO7B interacts with ANKS4B and USH1C through its first and second MyTH4-FERM (MF, myosin tail homology 4-band 4.1, ezrin, radixin, moesin) domains, respectively. ANKS4B interacts with MYO7B and USH1C, USH1C interacts with ANKS4B, MYO7B and CDHR2, and CDHR2 interacts with CDHR5 and USH1C (8,10–13). Whereas most of the binary interactions in the IMAC exhibit affinities in the range of ∼1-5 μM (8,12,13), the ANKS4B C-terminus binds to the USH1C N-terminal PDZ domain with a much higher affinity in the nM range (12,13). The extensive network of multivalent interactions among IMAC components may also lead to liquid-liquid phase separation upon complex formation (32).

Although most previous studies agree on the nature of interactions between different IMAC components (8,12,13), there are some points of conflict. Cell-based pulldown assays have demonstrated binding between CDHR5 and both USH1C and MYO7B (11). However, subsequent *in vitro* studies failed to detect binding between CDHR5 and USH1C (13), and the binding affinity between CDHR5 and MYO7B has not been measured. CDHR2 was shown to interact with the tail of MYO7B in pulldowns (11), although *in vitro* binding was measured to be ∼100-fold weaker than other IMAC interactions (13). Thus, how CDHR5 links to MYO7B and the underlying actin cytoskeleton, as well as the significance of direct binding between MYO7B and CDHR2 remain open questions. Biochemical evidence also indicates that three cytoplasmic components of the IMAC (MYO7B, ANKS4B, USH1C) assemble into a tripartite complex with 1:1:1 stoichiometry (11–13). Although this complex has been proposed to serve as a transport complex responsible for delivering protocadherins to the tips of microvilli (9,12), direct evidence for this model is lacking.

The complexity of the IMAC has made it challenging to study the role of individual binary interactions in complex formation and function in cells. One strategy relies on a KD/rescue approach, where atomic structure-guided mutagenesis is used to create IMAC component mutants that have impaired interactions with a specific binding partner; these mutants are then tested for their ability to rescue their own KD phenotype (e.g. defective BB assembly and microvillar clustering). Although this approach has been successful (8,11), the loss-of-function models needed to perform such a dissection have also proven difficult to generate due to perturbations in KD cell physiology. For example, CACO-2_BBE_ cells lacking USH1C expression exhibit a significantly reduced growth rate (unpublished observations). One alternative to a traditional KD/rescue approach is to reconstitute specific IMAC interactions in cells that do not normally express IMAC proteins. Along these lines, we developed a heterologous system that takes advantage of HeLa cells to study individual IMAC interactions and their resulting impact on the dynamics and morphology of surface protrusions, in this case filopodia. Our approach is based on a chimeric myosin that consists of the motor domain of MYO10 fused to the cargo binding tail domain of MYO7B. This synthetic myosin demonstrates robust targeting to the distal tips of HeLa cell filopodia and was able to drive enrichment of all other individual IMAC components - USH1C, ANKS4B, CDHR2 and CDHR5 - to these sites. Strikingly, we also observed that CDHR2 tip enrichment leads to “inter-filopodial adhesion”, which is coupled to a dynamic tip alignment activity that promotes length matching of adjacent protrusions. Our findings reveal that individual binary interactions between IMAC components and the MYO7B tail are robust enough to support transport and tip accumulation, and that the formation of a full tripartite complex is not necessary for the enrichment of microvillar protocadherins at the distal tips. These results also provide a mechanistic framework for understanding how BB microvilli achieve their characteristic length uniformity during enterocyte differentiation.

## RESULTS

### A heterologous in-cell assay for studying IMAC component interactions

To establish a system for studying IMAC component interactions and function, we focused on the filopodia that extend from the surface of HeLa cells. These finger-like protrusions are similar in size and shape to epithelial microvilli and, given that they extend parallel to the X-Y image plane, are also amenable to high temporal and spatial resolution imaging (Fig. 1B). However, full-length MYO7B, which is natively expressed in polarized epithelial cells and exhibits slow ATPase kinetics (33,34), does not efficiently translocate to the distal tips of filopodia (9), most likely because of the robust retrograde flow of the supporting core bundles in these structures. To circumvent this kinetic limitation, we created a chimeric myosin based on the filopodial motor, myosin-10 (MYO10); this approach was inspired by previous studies that took advantage of MYO10 motor domain (MD) tip targeting potential in a novel protein-protein interaction assay (35). Although full-length MYO10 actively transports numerous molecular cargoes to the distal tips of filopodia (36–38), the minimal domains necessary for distal tip targeting include the motor domain, IQ motifs, stable single a-helix (SAH), and anti-parallel coiled coil (CC). A truncated construct containing only these domains lacks cargo binding potential and exhibits robust tip targeting in HeLa cells (MY010MD, Fig. 1C,D). We generated the chimeric myosin by fusing the MYO7B cargo-binding tail domain to the C-terminus of MYO10MD (MYO10-MYO7B, Fig. 1C). Our expectation was that this chimeric motor protein would exhibit the motile properties and filopodial tip targeting of MYO10, while maintaining the cargo-binding potential of MYO7B. To confirm tip targeting, we expressed N-terminally EGFP-tagged MYO10-MYO7B in HeLa cells. Similar to EGFP-MYO10MD, EGFP-MYO10-MYO7B showed robust targeting to the distal tips of filopodia (Fig. 1E), indicating its utility for analyzing interactions with other IMAC components.

### The MYO10-MYO7B chimera transports IMAC scaffolding proteins to filopodia tips

We first sought to determine if MYO10-MYO7B was capable of transporting individual IMAC components to filopodial tips. ANKS4B and USH1C are modular scaffolding proteins with well-characterized binding interfaces on MYO7B MF1 and MF2, respectively (Fig. 1A) (8,12,13). To determine if binding to the MYO7B tail was sufficient for transporting these scaffolds tipward in the absence of other complex components, we transfected HeLa cells with mCherry-tagged ANKS4B or USH1C along with either EGFP-tagged MYO10MD or MYO10-MYO7B constructs. When expressed with the MYO10MD negative control, neither ANKS4B nor USH1C showed enrichment at the tips of filopodia (Fig. 2A,C). In contrast, co-expression with MYO10-MYO7B resulted in robust co-localization and tip targeting of both ANKS4B and USH1C (Fig. 2B,D). We quantified these results by generating ratios of the fluorescence intensity at the filopodial tip relative to the base; here, a purely soluble protein would yield a ratio of ∼1, whereas a protein highly enriched at the tips would generate a ratio of > 1. Using this quantification, both ANKS4B and USH1C showed ∼3-fold enrichment at the distal tips when expressed with MYO10-MYO7B relative to MYO10MD (Fig. 2E,F). Next we determined if the observed enrichment of scaffolding proteins at filopodial tips was a result of directed transport by the MYO10-MYO7B chimera. Live cell total internal reflection fluorescence microscopy (TIRFM) revealed the co-translocation of USH1C and MYO10-MYO7B along filopodial protrusions (Fig. 2G, Supplemental Movie 1). Large colocalized puncta of USH1C and MYO10-MYO7B moved retrograde from filopodial tips back toward the cell body (arrowheads, Fig. 2H) likely driven by actin core treadmilling (38), whereas smaller colocalized puncta demonstrated fast tipward movements (arrowheads, Fig. 2I) as expected for MYO10 driven transport (39). Interestingly, changes in MYO10-MYO7B puncta intensity were matched by parallel changes in USH1C intensity (Fig. 2J), suggesting that binding between the chimeric motor and its cargo is stoichiometric. The tight spatial coupling and intensity scaling of USH1C and MYO10-MYO7B puncta in these time-lapse data indicate that USH1C binding to the MYO7B tail domain, in the absence of other IMAC components, is stable enough to support its directed transport to the distal tips.

**Figure 2:**
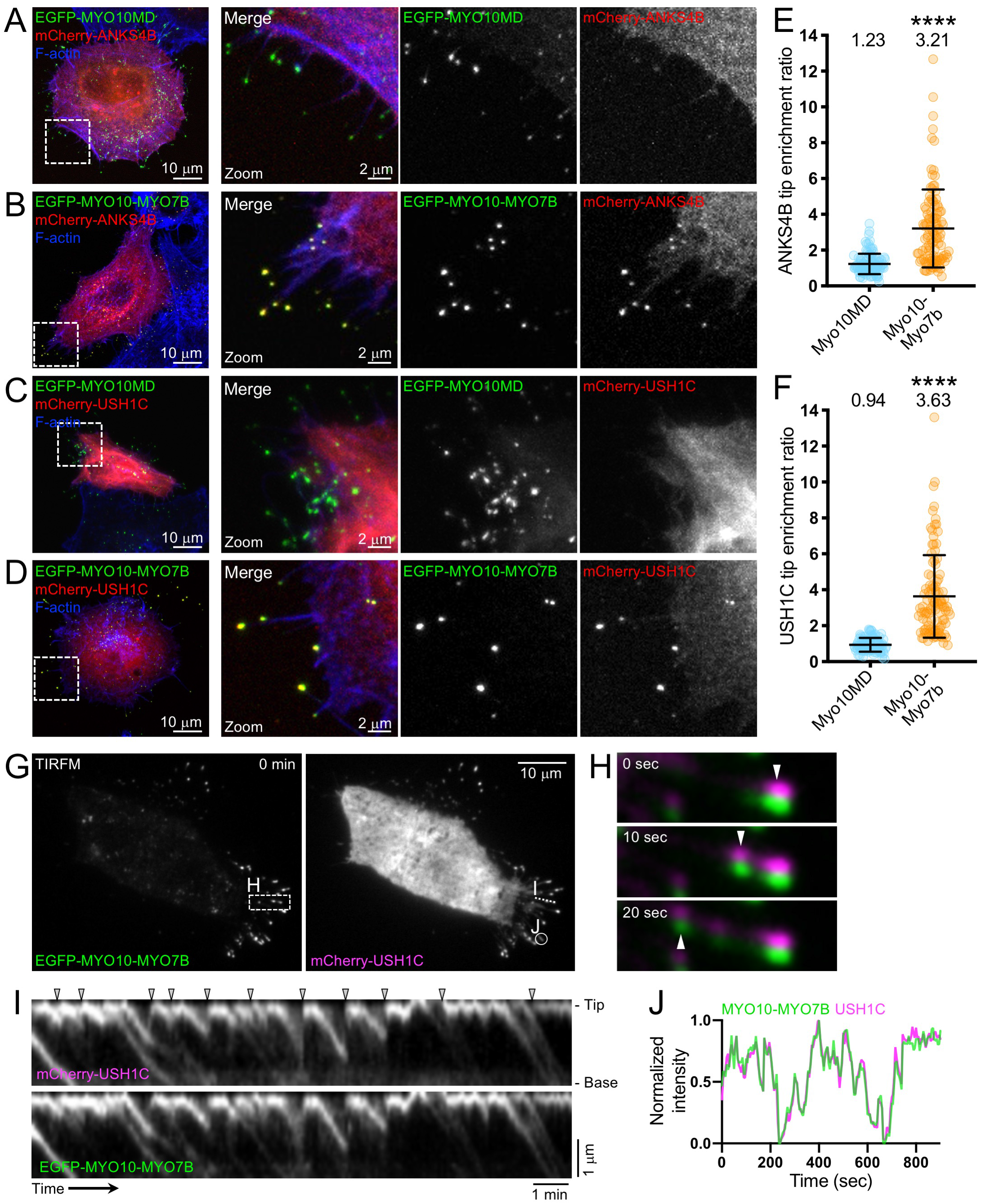
The MYO10-MYO7B chimeric myosin delivers scaffolding proteins ANKS4B and USH1C to the tips of filopodia. Confocal images of HeLa cells co-expressing: (A) EGFP-MYO10MD (green) and mCherry-ANKS4B (red), (B) EGFP-MYO10-MYO7B (green) and mCherry-ANKS4B (red), (C) EGFP-MYO10MD (green) and mCherry-USH1C (red), or (D) EGFP-MYO10-MYO7B (green) and mCherry-USH1C (red). In A-D, all cells are counterstained with phalloidin to label F-actin (blue); the white dashed outlined box in the overview panel at left indicates the region shown in the merge and single channel zoom panels to the right. (E) Quantification of ANKS4B filopodial tip enrichment ratios in cells expressing MYO10MD (n = 107 filopodia from 9 cells) or MYO10-MYO7B (109 filopodia from 11 cells); ****p < 0.0001. (F) Quantification of USH1C filopodial tip enrichment ratios in cells expressing MYO10MD (n = 86 filopodia from 7 cells) or MYO10-MYO7B (173 filopodia from 10 cells); ****p < 0.0001. For E and F, each data point represents the enrichment ratio from a single filopodium, and population means are indicated at the top of each scatter plot. (G) TIRFM images of a live HeLa cell co-expressing EGFP-MYO10-MYO7B (green, left) and mCherry-USH1C (magenta, right). (H) Vertically montaged images of 0, 10, and 20 sec timepoints from the image series of the cell shown in G, taken from the outlined box marked H. EGFP-MYO10-MYO7B (green) and mCherry-USH1C (magenta) are offset vertically to facilitate visualization of the simultaneous movement of both signals; white arrowheads mark puncta that are moving retrograde back to the cell body. (I) Kymographs of the mCherry-USH1C (top) EGFP-MYO10-MYO7B (bottom) signals generated from the cell shown in G, taken from the filopodium marked I. Tip and base of the filopodium are marked with B and T to the right; gray arrows mark rapid tip-ward transport events. (J) Plot of EGFP-MYO10-MYO7B (green) and mCherry-USH1C (magenta) fluorescence intensities over time from the cell shown in G, taken from the region of interest marked J. Dimensions of scale bars are indicated in figure panels.

### The MYO10-MYO7B chimera transports CDHR2s to the distal tips

*In vitro* biochemical studies revealed that the CDHR2 cytoplasmic domain (CD) binds weakly to the tail of MYO7B, with an affinity of ∼140 μM (13), although the structural details of this interaction and the motifs involved are unknown. In contrast, the CDHR2 C-terminal PDZ-binding motif (PBM) interacts with the second PDZ domain of USH1C with a much higher affinity of ∼1.4 μM (13). Given that USH1C binding to CDHR2 is ∼100-fold stronger than binding to MYO7B, the physiological significance of weak CDHR2/MYO7B binding remains unclear. These differential affinities might also suggest that CDHR2 relies on direct binding to USH1C and thus, indirect binding to MYO7B, for its delivery to the distal tips of microvilli. From this perspective, we tested the CDHR2/MYO7B interaction using the in-cell assay introduced above. Because CDHR2 is a transmembrane protein, expression of an EGFP-tagged construct resulted in signal along the length of filopodia when expressed in HeLa cells alone or with the MYO10MD control (Fig. 3A,B). Somewhat unexpectedly, however, co-expression of CDHR2-EGFP with the MYO10-MYO7B chimeric motor resulted in strong distal tip enrichment (Fig. 3C,E). CDHR2 targeting in this context was also dependent on its CD, as deletion of this region resulted in a loss of tip enrichment in the presence of MYO10-MYO7B (Fig. 3D,E). Importantly, and similar to what we observed with USH1C, live TIRFM showed that CDHR2 accumulation at the distal tips was the result of robust co-translocation with the MYO10-MYO7B chimera (Supplemental Movie 2).

**Figure 3:**
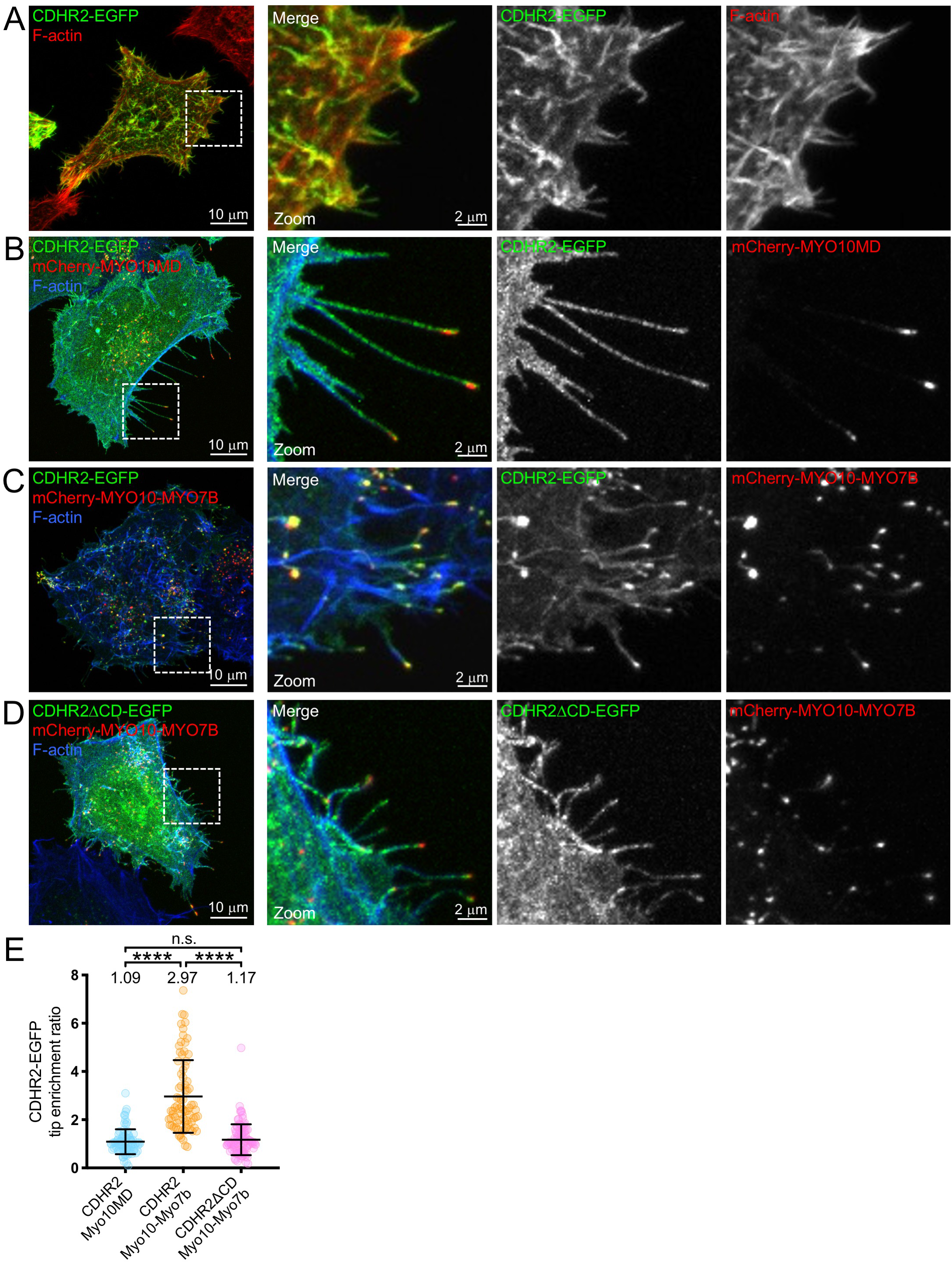
The MYO10-MYO7B chimeric myosin delivers CDHR2 to the tips of filopodia. Confocal images of HeLa cells expressing: (A) CDHR2-EGFP (green), (B) CDHR2-EGFP (green) and mCherry-MYO10MD (red), (C) CDHR2-EGFP (green) and mCherry-MYO10-MYO7B (red), or (D) CDHR2DCD-EGFP (green) and mCherry-MYO10-MYO7B (red). In A-D, cells are counterstained with phalloidin to label F-actin (red in A, blue in B-D); the white dashed outlined box in the overview panel at left indicates the region shown in the merge and single channel zoom panels to the right. Dimensions of scale bars are indicated in figure panels. (E) Quantification of filopodial tip enrichment ratios in cells co-expressing CDHR2 + MYO10MD (n = 78 filopodia from 7 cells), CDHR2 + MYO10-MYO7B (79 filopodia from 5 cells), or CDHR2DCD + MYO10-MYO7B (104 filopodia from 8 cells), ****p < 0.0001. In E, each data point represents the enrichment ratio from a single filopodium, and population means are indicated at the top of each scatter plot.

Although CDHR2 tip enrichment and co-translocation with MY010-MYO7B might reflect a direct interaction, we first explored the possibility that other IMAC proteins are endogenously expressed in HeLa cells, where they might facilitate CDHR2 tip accumulation. SDS-PAGE/western blot analysis revealed that the only detectable IMAC protein in HeLa whole cell lysates is USH1C; moreover, we were able to detect USH1C in a wide range of cell culture models (Supplemental Fig. 1). To determine if endogenous USH1C couples CDHR2 to MYO10-MYO7B in our HeLa cell tip targeting assay, we introduced a charge reversal point mutation, K1918E, into MYO10-MYO7B MF2 to abolish USH1C binding (8). The K1918E mutation was designed based on a crystal structure of the USH1C-PDZ3/MYO7B-MF2 complex and its ability to disrupt this binary interaction has been validated *in vitro* and in cells (8). We first confirmed that MYO10-MYO7B-K1918E was unable to promote the enrichment of exogenously expressed USH1C in our system (Fig. 4A). Next, we co-expressed MYO10-MYO7B-K1918E with CDHR2 and found that both constructs still exhibited strong enrichment and co-localization at the distal tips of filopodia (Fig. 4B-D). Interestingly, quantification of enrichment ratios revealed that the K1918E mutation reduced CDHR2 tip targeting slightly, but tip enrichment was still significantly greater than when co-expressed with the MYO10MD negative control (Fig. 4E). Taken together, we conclude that direct interactions with the MYO7B tail domain support the delivery of CDHR2 to the distal tips of protrusions, and that this activity may be modestly enhanced in the presence of USH1C.

**Figure 4:**
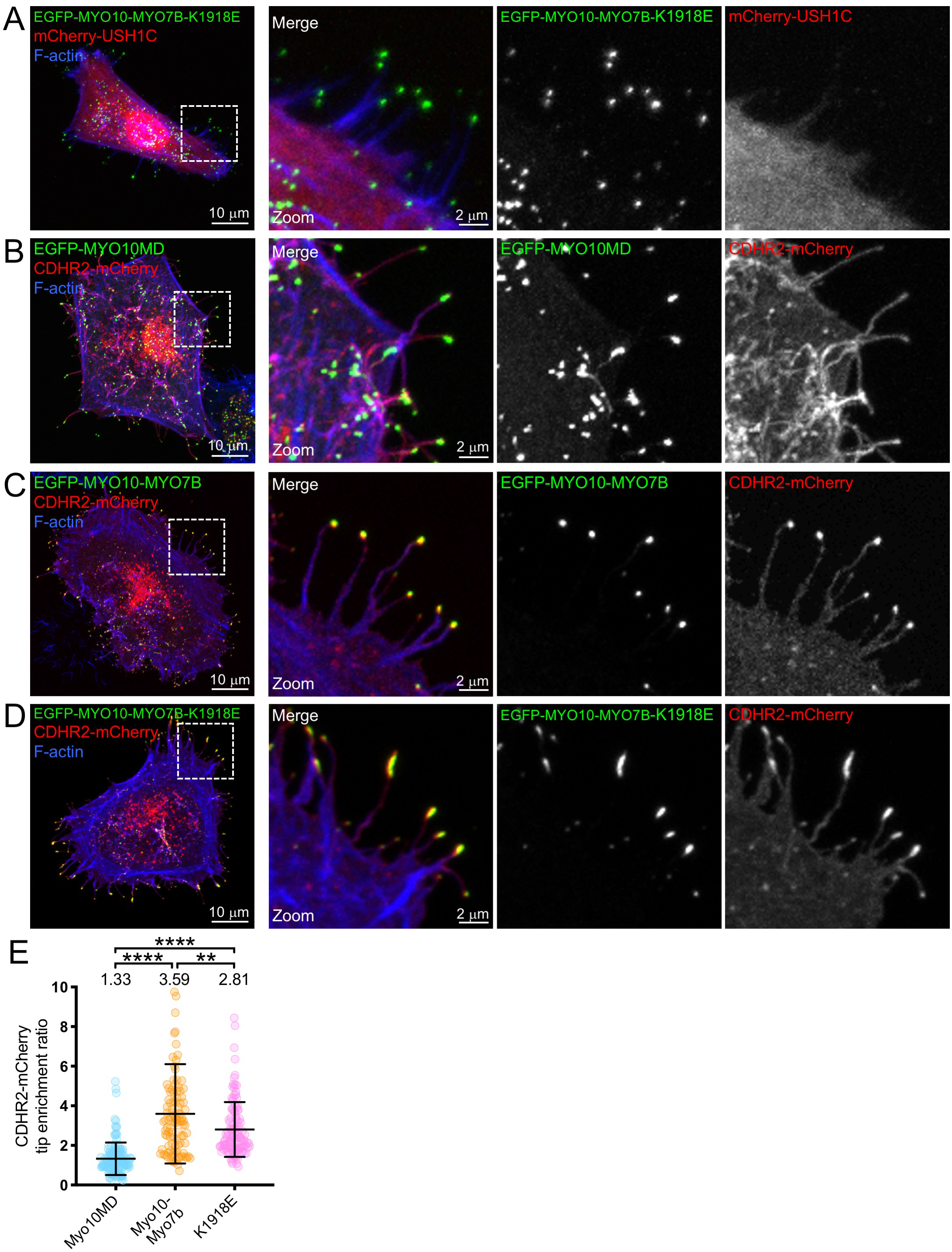
CDHR2 tip delivery does not depend on USH1C binding to the MYO10-MYO7B chimera. Confocal images of HeLa cells expressing: (A) EGFP-MYO10-MYO7B-K1918E (green) and mCherry-USH1C (red), (B) EGFP-MYO10MD (green) and CDHR2-mCherry (red), (C) EGFP-MYO10-MYO7B (green) and CDHR2-mCherry (red), or (D) EGFP-MYO10-MYO7B-K1918E (green) and CDHR2-mCherry (red). In A-D, cells are counterstained with phalloidin to label F-actin (blue); the white dashed outlined box in the overview panel at left indicates the region shown in the merge and single channel zoom panels to the right. Dimensions of scale bars are indicated in figure panels. (E) Quantification of CDHR2 filopodial tip enrichment ratios in cells co-expressing MYO10MD (n = 135 filopodia from 7 cells), MYO10-MYO7B (102 filopodia from 7 cells), or MYO10-MYO7B-K1918E (118 filopodia from 7 cells); ****p < 0.0001. In E, each data point represents the enrichment ratio from a single filopodium, and population means are indicated at the top of each scatter plot.

### The MYO10-MYO7B chimera promotes distal tip enrichment of CDHR5

We next sought to determine if the MYO10-MYO7B chimera promotes the distal tip enrichment of CDHR5. CDHR5 trafficking to the plasma membrane in HeLa cells is inefficient, but a fraction of CDHR5 clearly localized to the plasma membrane (Fig. 5A). Similar to other IMAC components, when coexpressed with the MYO10-MYO7B chimeric motor, CDHR5 exhibited enrichment at the distal tips of filopodia (Fig. 5C,E). To explore a potential role for USH1C in CDHR5 localization, we again took advantage of the MYO10-MYO7B-K1918E mutant chimera. Although cell-based pulldowns showed that CDHR5 binds to USH1C in a manner similar to CDHR2 (11), *in vitro* studies were unable to detect binding (13). Nevertheless, similar to our results with CDHR2 (Fig. 4), the K1918E mutation only slightly decreased CDHR5 tip targeting, and the enrichment ratio was still significantly increased compared to the MYO 10MD negative control (Fig. 5B-E). As with CDHR2, we conclude that direct binding to MYO7B drives the tip localization of CDHR5, and that USH1C may stabilize MYO7B/CDHR5 interactions.

**Figure 5:**
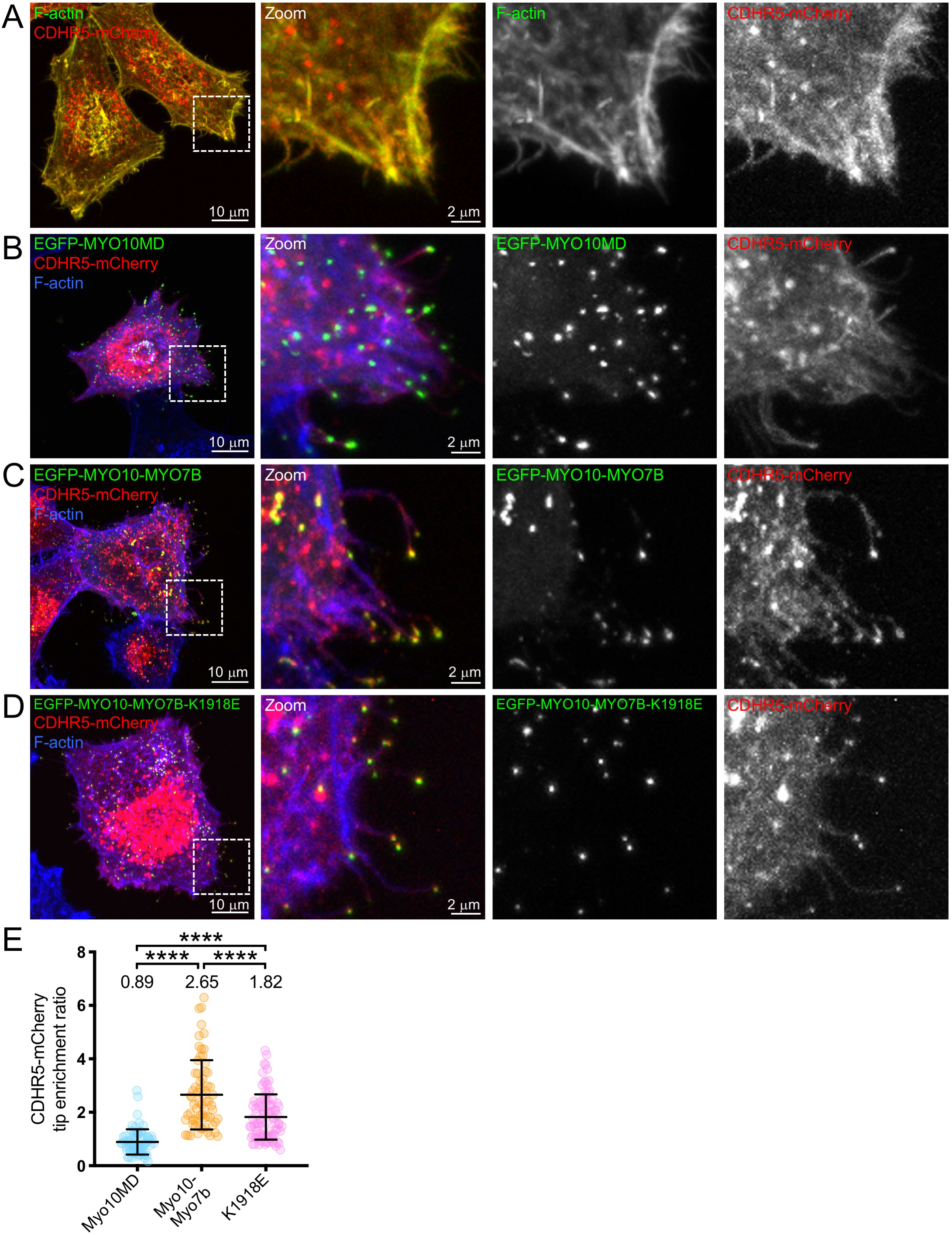
The MYO10-MYO7B chimeric myosin delivers CDHR5 to the tips of filopodia. Confocal images of HeLa cells expressing: (A) CDHR5-mCherry (red), (B) CDHR5-mCherry (red) and EGFP-MYO10MD (green), (C) CDHR5-mCherry (red) and EGFP-MYO10-MYO7B (red), or (D) CDHR5-mCherry (red) and EGFP-MYO10-MYO7B-K1918E (green). In A-D, cells are counterstained with phalloidin to label F-actin (green in A, blue in B-D); the white dashed outlined box in the overview panel at left indicates the region shown in the merge and single channel zoom panels to the right. Dimensions of scale bars are indicated in figure panels. (E) Quantification of CDHR5 filopodial tip enrichment ratios in cells co-expressing MYO10MD (n = 71 filopodia from 12 cells), MYO10-MYO7B (66 filopodia from 5 cells), or MYO10-MYO7B-K1918E (94 filopodia from 7 cells), ****p < 0.0001. In E, each data point represents the enrichment ratio from a single filopodium, and population means are indicated at the top of each scatter plot.

### Tip enriched CDHR2 drives inter-filopodial adhesion and dynamic length matching

Our previous studies established that the CDHR2 ectodomain binds weakly to itself (11), although the biological role of such homophilic adhesion relative to the stronger heterophilic binding with CDHR5 remains unclear. Thus, we took advantage of the fact that we could target exogenous CDHR2 to filopodial tips by simply co-expressing the MYO10-MYO7B chimera, to examine the specific impact of homophilic adhesion on the morphology and dynamics of filopodial protrusions. To better visualize filopodia on the dorsal surface of HeLa cells, we turned to super-resolution structured illumination microscopy (SIM). Relative to control cells (Fig. 6A-C), cells expressing CDHR2 and MYO10-MYO7B presented striking examples of contact between the tips of neighboring filopodia, which corresponded to regions where CDHR2 levels were high (arrows, Fig. 6D). These structures were strikingly reminiscent of tipi-like clusters of microvilli found on the surface of CACO-2_BBE_ cells early in differentiation (11). In some CDHR2/MYO10-MYO7B co-expressing cells, we also observed protrusion bundles consisting of multiple filopodia (Fig. 6E), where most of the distal tips were clustered together at a similar distance from the cell surface. Using lateral SIM views, we scored the frequency of filopodia engaged in tip-to-tip adhesion on the surface of negative controls cells (Fig. 6A-C) and cells co-expressing CDHR2 and MYO10-MYO7B (Fig. 6D). Although dorsal filopodia were found on the surface of cells expressing either CDHR2 or MYO10-MYO7B alone, we observed significantly more tip-to-tip contacts between adjacent filopodia on cells expressing both constructs (Fig. 6F). Examination of negative control and CDHR2/MYO10-MYO7B co-expressing cells using scanning EM confirmed this point (Fig. 6G,H) and additionally, revealed several striking examples of length-matched adherent filopodia (Fig. 6H, panels i-v). Together, these findings suggest that the elongation of a filopodium may be limited when its distal tip forms adherent attachments to adjacent protrusions.

**Figure 6:**
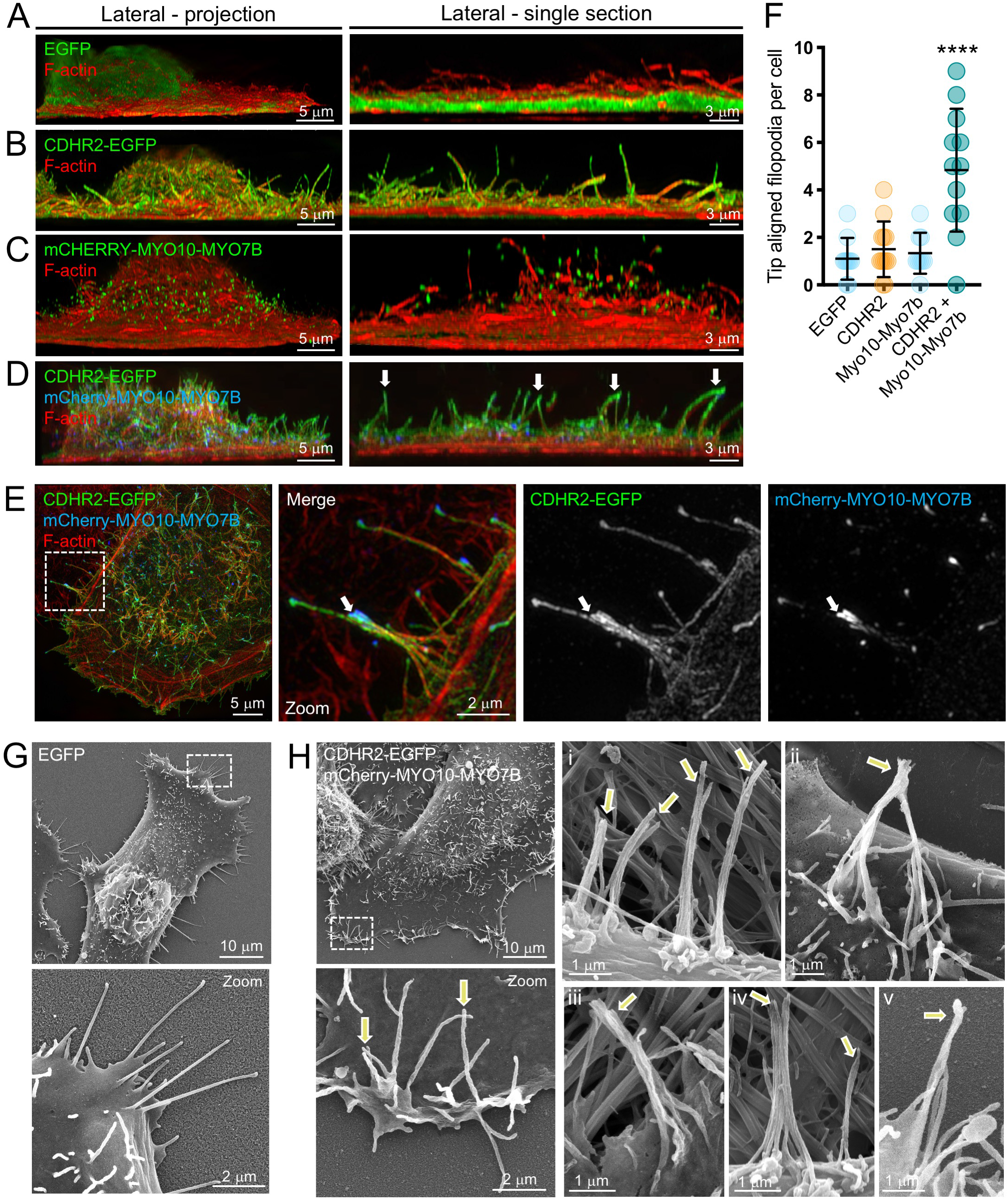
Tip targeted CDHR2 promotes inter-filopodial adhesion and spatial alignment of distal tips. Lateral views of super-resolution SIM volumes of HeLa cells expressing (A) EGFP, (B) CDHR2-EGFP, (C) mCherry-MYO10-MYO7B, or (D) CDHR2-EGFP and mCherry-MYO10-MYO7B. *Left*, lateral view of the full volume projection; *right*, lateral view of a single representative section from the volume. In A-D, cells are counterstained with phalloidin to label F-actin (red). In D, white arrows indicate tip-tip adhesions formed between adjacent filopodia. (E) Super-resolution SIM *en face* volume projection of a HeLa cell expressing CDHR2-EGFP (green) and mCherry-MYO10-MYO7B (blue), and counterstained with phalloidin to label F-actin (red). The white dashed outlined box in the overview panel at left indicates the region shown in the merge and single channel zoom panels to the right. In the zoom panels in E, white arrows highlight a large filopodial bundle consisting of multiple protrusions. (F) Scoring the frequency of tip-aligned dorsal filopodia on the surface of HeLa cells expressing individual control constructs (EGFP, CDHR2-EGFP, mCherry-MYO10-MYO7B) or co-expressing CDHR2-EGFP and mCherry-MYO10-MYO7B, as visualized in lateral views of SIM image volumes; each data point corresponds to the value scored from a single cell, ****p < 0.0001. (G) Scanning EM images of control HeLa cells expressing EGFP only. The white dashed outlined box in the top overview panel indicates the region shown in the bottom zoom panel. (H) Scanning EM images of HeLa cells expressing CDHR2-EGFP and mCherry-MYO10-MYO7B. The white dashed outlined box in the top overview panel indicates the region shown in the bottom zoom panel; panels (i-v) to the right show additional examples of adherent and length-matched filipodia. Dimensions of scale bars are indicated in figure panels.

To further explore the mechanism of length matching between adherent filopodia, we used TIRFM to visualize protrusion dynamics on the surface of live CDHR2/MYO10-MYO7B co-expressing HeLa cells relative to EGFP expressing controls. Because TIRFM confines fluorescence excitation to the coverslip surface, these imaging experiments focused on surface-associated filopodia. Live cell TIRFM revealed numerous examples of filopodia bending or leaning toward neighboring protrusions in CDHR2/MYO10-MYO7B co-expressing cells, relative to negative control cells where filopodia were generally straighter and non-adherent (Fig. 7A-C; Supplemental Movie 3). We also observed clusters of filopodia with clear tip-to-tip contacts (Fig. 7B, outlined white arrowheads). Moreover, in some cases we were able to capture the dynamics that gave rise to these clusters. In watching these events play out over time, we found that the tip of a nascent filopodium first attached to a pre-existing adjacent protrusion, usually somewhere along its length (Fig. 7B, solid white arrowhead; Supplemental Movie 3). The newer filopodium then elongated while its tip ‘slid’ towards the distal end of the pre-existing structure (Fig. 7C, white arrow; Supplemental Movie 3). Once the newer tip reached the position of the older tip, the structures appeared to stabilize. To determine if this dynamic tip alignment activity was specific to the surface-association of filopodia visualized using TIRFM or instead represented a general behavior of all filopodia with tip-enriched CDHR2, we performed similar imaging studies using spinning disk confocal microscopy, which allowed volume acquisition and thus visualization of dorsal (i.e. non-surface associated) filopodia. Confocal timelapse data revealed clear examples of upright dorsal filopodia colliding and adhering with adjacent protrusions, followed by tip sliding and alignment as observed in TIRFM datasets (Fig. 7D, numbered solid white arrowheads indicate distinct filopodia; Supplemental Movie 4). These observations indicate that distal tip adhesion complexes form through a dynamic tip sliding and spatial alignment mechanism, which ultimately constrains the length of all protrusions in the adherent cluster.

**Figure 7:**
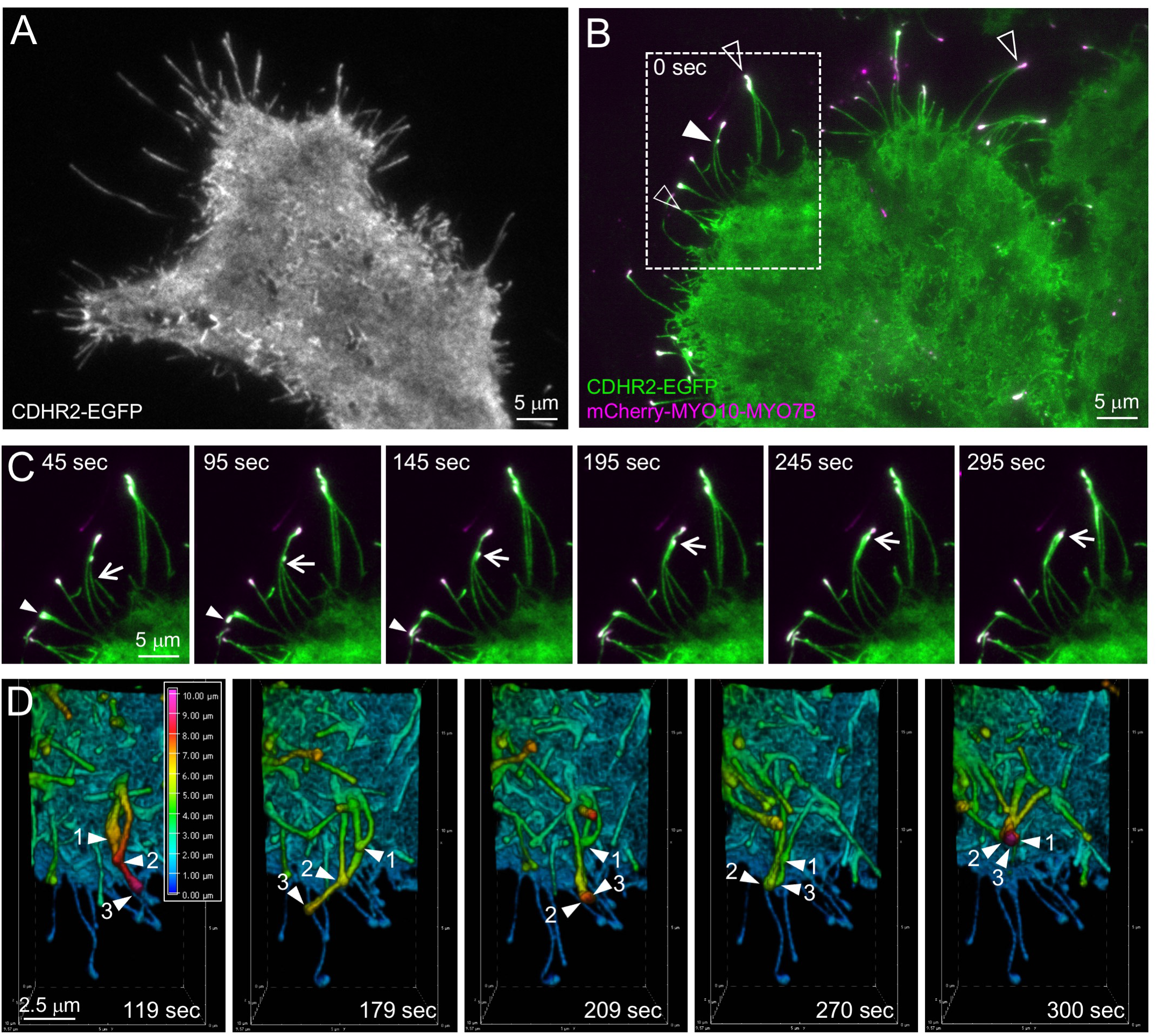
New filopodia with tip targeted CDHR2 collide with neighboring protrusions and exhibit dynamic length matching. (A) Live TIRFM image of a HeLa cell expressing CDHR2-EGFP alone. (B) Live TIRFM image of a HeLa cell expressing CDHR2-EGFP (green) and mCherry-MYO10-MYO7B (magenta). The white outlined arrowheads indicate tip-tip adhesions formed between adjacent filopodia that are roughly matched in length; the white solid arrowhead highlights a case where the distal tip of a new filopodium has bound along the shaft of a pre-existing neighboring protrusion. (C) Time-lapse montage of the white outlined boxed region highlighted in B; white arrow marks the changing position of the distal tip of the new filopodium as it elongates, sliding along the shaft of the pre-existing neighboring protrusion. (D) Live spinning disk confocal time-lapse montage of a HeLa cell expressing CDHR2-EGFP and mCherry-MYO10-MYO7B; a 3D reconstructed and depth-coded version of the CDHR2-EGFP signal is shown here. Dorsal filopodia which are highly dynamic and remain detached from the coverslip surface, also demonstrate robust length matching following the formation of adherent contacts; numbered arrowheads in this example highlights three distinct filopodia, which align their distal tips over time. Dimensions of scale bars are indicated in figure panels.

## DISCUSSION

Localization studies in native tissues show that all IMAC components demonstrate robust enrichment at the tips of microvilli in kidney and intestinal epithelial cells (7,9–11,14), and loss-of-function investigations have confirmed that tip accumulation is critical for the function of the IMAC in microvillar packing and brush border maturation (7,9–11,14). Previous cell biological studies also indicate that MYO7B plays an important role in this localization, although the precise mechanism has remained somewhat ambiguous (9). MYO7B binds to all other IMAC components (10,11), demonstrates catalytic and motor activity (33), and mutations in its motor domain impair localization and tip enrichment of associated IMAC cargoes (9). Moreover, overexpressed EGFP-MYO7B accumulates at the tips of microvilli in differentiating epithelial cells (9), and in studies of other motor proteins, such tip accumulation is usually regarded as a telltale sign of active, end-directed transport (38,40). Taken together, all of this evidence points to a model where MYO7B actively transports IMAC components, either individually or in complex, to the distal tips of microvilli.

Accepting a role for MYO7B in active transport, however, remains difficult as several questions remain unanswered and key pieces of supporting data are lacking. For example, how MYO7B navigates the extremely tight confines of the intramicrovillar cytoplasm and overcomes the steric resistance sure to be encountered during tipward transport remains unclear. There is also the question of how effective a motor protein would be at sliding a transmembrane cargo (e.g. CDHR2) through the plane of the plasma membrane, on its way from the protrusion base to tip. This problem becomes even more significant if *trans* adhesion complexes form between adjacent microvilli first and are then subsequently transported tipward. Moreover, clear time-lapse imaging evidence showing the movement of MYO7B bound to its IMAC cargoes within microvilli has remained elusive. This is most likely due to several technical reasons, including the vertical orientation of microvilli. Apical microvilli extend through multiple z planes and given the low resolution of the z-axis on optical microscopes, capturing movement of MYO7B (with or without IMAC cargoes) in these structures has proven difficult. In the absence of this data, competing hypotheses remain in consideration. For example, MYO7B might use some combination of slow motility and diffusion to arrive at the tips, where in turn, it serves as a point of attachment for other IMAC proteins and promotes their passive accumulation at these sites. This would be consistent with previous data showing that a mutant variant of MYO7B unable to generate force or motion was able to weakly target to microvillar tips and at least partially rescue the tip enrichment of CDHR2 in the CACO-2_BBE_ model system (9).

Here we describe an approach for studying the ability of individual IMAC cargoes to interact with the MYO7B tail domain and additionally, determine whether these interactions are robust enough to support component delivery to and enrichment at the distal tips of actin-based protrusions. Based on the “nanoscale pulldown” approach originally described by Bird and colleagues (35), our assay takes advantage of a chimeric myosin motor consisting of the MYO10 motor domain coupled to the MYO7B tail domain; the MYO10 motor permits entry into and transport to the tips of HeLa cell filopodia, whereas the MYO7B tail allows us to examine interactions with individual IMAC cargoes. Importantly, because most IMAC components are not expressed at detectable levels in HeLa cells, heterologous expression allows us to isolate and study specific binary interactions by simply co-expressing the MYO10-MYO7B chimera with the cargo of interest. Using this system, we found that the chimeric motor is capable of driving the tip enrichment of all MYO7B IMAC binding partners: CDHR2, CDHR5, ANKS4B, and USH1C. Moreover, live imaging studies clearly showed that IMAC cargoes bind to and move with the chimeric motor. Thus, individual binary interactions with the MYO7B tail domain are robust enough to endure active transport and delivery to the distal tips. Collectively, our new findings underscore the potential for MYO7B to support directed transport of its cargoes to the distal tips of microvilli and provide further insight on mechanisms of IMAC accumulation at these sites.

Unexpectedly, we found that even in the absence of interactions with scaffolding factors ANKS4B or USH1C, binding interactions between MYO7B and the microvillar protocadherins CDHR2 and CDHR5 were robust enough to support filopodial tip targeting in the heterologous HeLa cell system. Although the in-cell nature of this assay makes it difficult to state with certainty that the MYO7B/protocadherin interactions observed here represent direct binding, we suspect this is the case for two reasons. First, whereas we were able to detect USH1C in HeLa cell lysates, a mutation in the MYO7B tail that abolishes USH1C binding had little impact on the ability of the MYO10-MYO7B chimera to deliver CDHR2 or CDHR5 to the distal tips. Moreover, direct binding between the MYO7B tail and CDHR2 is consistent with previous pulldown studies that demonstrated direct binding albeit with low affinity *in vitro* (11,13). We suspect the limitations of low binding affinity might be diminished in the tight confines of the intra-filopodial cytoplasm.

Despite the potential for direct binding between MYO7B and the microvillar protocadherins, other experiments indicate that ANKS4B and USH1C are required for normal complex localization and function in CACO-2_BBE_ intestinal epithelial cells and *in vivo* (10,11). This might suggest that, in addition to simply linking CDHR2 or CDHR5 to MYO7B, the scaffolding proteins have additional functions such as motor activation or spatial/temporal regulation of IMAC formation. Indeed, biochemical evidence suggests that the MYO7B tail domain exists in an auto-inhibited state that prevents interaction with ANKS4B and potentially other IMAC components (10). In this scheme, USH1C binding to the MYO7B tail releases autoinhibition and subsequently enables interaction with ANKS4B; MYO7B/USH1C binding is not dependent on the interaction between ANKS4B and USH1C. The MYO10-MYO7B chimeric myosin used in the current studies is not expected to be subject to strong autoinhibitory regulation.

Given that the MYO10-MYO7B chimera drives CDHR2 accumulation at the tips of filopodia, we took advantage of this finding to examine the impact of tip enriched adhesion on the dynamics and morphology of filopodia. Previous studies established that CDHR2 is capable of interacting with itself to form *trans* homophilic complexes (11). Indeed, in fixed and live cell observations, we found clear evidence of tip-tip adhesion complex formation, with adjacent filopodia leaning toward each other, making physical contact at their tips. In time-lapse observations, we also observed a striking tip alignment activity, where the tip of an elongating nascent protrusion first attached to a pre-existing adjacent filopodium along its length (Fig. 7, Supplemental Movies 3 and 4). As the newer filopodium elongated, its distal tip slid towards the tip of the pre-existing filopodium. Once the newer tip reached the pre-existing tip, the two ends formed a stable adhesive structure that was relatively long-lived. Stable interactions form between the tips, as these are the sites along the protrusion axis where the density of CDHR2 molecules is highest. We also noted that the adherent clustering of distal tips in this manner seemed to constrain the lengths of adjacent filopodia, which were commonly similar in length (Fig. 6). In this light, tip-tip adhesion might represent a length matching mechanism for adjacent protrusions; the distal tip of the pre-existing protrusion could effectively serve as a barrier to the elongation of nascent protrusions once the ends become complexed together due to CDHR2-based adhesion. This is an intriguing idea given that length uniformity is one of the defining morphological features of the brush border microvilli (5). This hypothesis is also supported by recent studies of the CDHR2 KO mice, which revealed increased variability in microvillar length, splaying of protrusions, and a ∼30% decrease in the number of microvilli (7).

Although the *relative* lengths of adjacent protrusions might be constrained or matched through protocadherin-based tip-tip adhesion, other mechanisms likely control the *absolute* length of microvilli *in vivo*. Size sensing could occur directly through a dedicated reporter/ruler molecule or indirectly, through measurements of functional capacity (41). In the absence of direct size sensing, one mechanism recently highlighted for actin-based structures is linked to the abundance of precursors, in this case actin monomers. Indeed, recent studies suggest that G-actin is a limiting resource and that competition for subunits between actin nucleators and assembly factors (e.g. Arp2/3 and formins) dictates the size and architecture of the resulting F-actin array (42–45). Recent work from our group suggest that such a mechanism may be operational in the intestinal brush border (46). Future studies must focus on clarifying mechanisms of and interplay between absolute and relative size control for epithelial microvilli and related structures.

## EXPERIMENTAL PROCEDURES

### Construct cloning

The *Hs* cDNA construct of MYO7B used in this study corresponds to NCBI GI: 122937511, whereas the full-length *Bt* MYO10 construct was a kind gift from Dr. Richard Cheney. All synthetic constructs were generated by PCR and TOPO cloned into the pCR8 Gateway entry vector (Invitrogen). Restriction enzyme sites, point mutations, and refractory silent mutations were introduced using QuikChange site-directed mutagenesis (Agilent). All entry vectors were first verified by DNA sequencing and then shuttled into destination vectors pEGFP-C1 (Clontech) and pINDUCER20-EGFP-C1 (10), which were Gateway-adapted using the Gateway vector conversion kit (Invitrogen). The MYO10-MYO7B chimeric motor was constructed by fusing *Bt* MYO10 nt 1-2805 (encoding a.a. 1-935) in frame with *Hs* MYO7B nt 3021-6492 (encoding a.a. 964-2164) using Gibson assembly. All constructs were verified by DNA sequencing. All ANKS4B, USH1C, CDHR2, and CDHR5 constructs used in this study were described previously (10,11).

### Cell culture and transfections

All cells were cultured at 37°C and 5% CO_2_ in DMEM with high glucose and 2 mM L-glutamine supplemented with 10% FBS or with 20% FBS for CACO-2_BBE_ cells. Transfections were performed using Lipofectamine 2000 (Invitrogen) according to the manufacturer’s instructions and cells were allowed to recover overnight. The following day, cells were plated on glass-bottom dishes and/or coverslips coated with 25 μg/ml laminin (Sigma) and allowed to adhere overnight before live imaging or fixing/staining.

### Western blot analysis of cultured cells

To create lysates for western blotting, HeLa cells were grown in a T75 flask until 90% confluent, whereas CACO-2_BBE_ cells were seeded into T25 flasks and grown until 14 days post confluency. Cells were washed once with warm PBS and recovered in 5 mL of PBS using a cell scraper. Cells were pelleted at low speed and resuspended in RIPA buffer containing 1 mM ATP, 1 mM Pefabloc SC protease inhibitor (Roche), and 1x cOmplete ULTRA protease inhibitor cocktail (Roche). Cells were lysed by needle aspiration and centrifuged at 16,000 x *g* for 5 min at 4° C. The soluble fraction was recovered, hot SDS sample buffer was added, and samples were boiled for 3 min. Samples were then separated on 4-12% NuPAGE Bis-Tris gel (Novex) and transferred in Towbin buffer, pH 8.3, to nitrocellulose at 15 V overnight. Membranes were blocked for 1 hr in 5% milk in PBS containing 0.1% Tween-20 (PBS-T), washed once with PBS-T, and then incubated with primary antibodies against MYO7B (1:100; Sigma #HPA039131), CDHR2 (1:100; Sigma #HPA012569), CDHR5 (1:500; Sigma #HPA009081), ANKS4B (1:200; Sigma #HPA043523), USH1C (1:250; Sigma #HPA027398), or GAPDH (1:1000; Cell Signaling #2118L) in 5% milk PBS-T for 2 hrs. Membranes were washed three times with PBS-T and incubated with goat or donkey anti-rabbit 800IRDye (1:10,000; Li-Cor) in 5% milk PBS-T for 1 hr. Membranes were washed four times with PBS-T before being imaged using a Li-Cor Odyssey infrared imaging system. Images of membrane scans were contrast enhanced and quantified using ImageJ (NIH).

### Light microscopy

HeLa cells were washed once with warm PBS and fixed with 4% paraformaldehyde in PBS for 15 min at 37° C. After fixation, cells were washed three times with PBS and permeabilized with 0.1% Triton X-100 in PBS for 15 min at room temperature (RT). Cells were then washed three times with PBS and blocked overnight with 5% BSA in PBS at 4°C. Cells were washed once with PBS and immunostaining was performed using anti-CDHR2 (1:75; Sigma #HPA012569), anti-CDHR5 (1:250; Sigma #HPA009081), anti-USH1C (1:70; Sigma #HPA027398), and anti-MYO7B (1:25; Sigma #HPA039131) at RT for 2 hours. Coverslips were then washed three times with PBS and incubated with AlexaFluor488 goat anti-rabbit or AlexaFluor647 goat anti-rabbit and AlexaFluor568 phalloidin (1:200; Life Technologies) diluted 1:200 in PBS for 1 hour at 37°C. Cells were washed four times with PBS and coverslips were mounted using ProLong Gold Antifade Mountant (Life Technologies). Cells were then washed three times with PBS and stained with AlexaFluor568 phalloidin (1:200) in PBS for 30 min at 37° C. Coverslips were washed and mounted as above. For super-resolution microscopy, cells were fixed and permeabilized as above. Cells were then washed three times with PBS and blocked overnight with 5% BSA in PBS at 4°C. Cells were washed once with PBS and stained with anti-GFP (1:800; Aves labs #GFP1020) or anti-GFP and anti-mCherry (1:500; Invitrogen #M11217) for 2 hours at RT. Coverslips were then washed three times with PBS and incubated with AlexaFluor488 goat anti-chicken (1:200; Life Technologies) and AlexaFluor568 phalloidin (1:200; Life Technologies) or AlexFluor488 goat anti-chicken, AlexaFluor568 goat anti-rat (1:200; Life Technologies), and AlexaFluor647 phalloidin (1:100, Life Technologies) diluted PBS for 1 hour at 37°C. Cells were washed four times with PBS and coverslips were mounted using ProLong Gold Antifade Mountant. Tissue sections and fixed cells were imaged using a Nikon A1R laser-scanning confocal microscope. TIRF live cell imaging was performed on a Nikon TiE inverted light microscope equipped with 488 and 561 excitation LASERS, a 100x/1.49 NA TIRF objective, and a Photometrics Evolve EM-CCD camera. Structure illumination microscopy was performed using an Applied Precision DeltaVision OMX (GE Healthcare) equipped with a 60x Plan-Apochromat N/1.42 NA oil immersion objective (Olympus) and processed using softWorx software (GE Healthcare). For live cell imaging, cells were maintained in a humid environment at RT and 5% CO_2_ using a stage-top incubation system (Tokai Hit). Image acquisition was controlled with Nikon Elements software.

### Electron microscopy

All SEM reagents were purchased from Electron Microscopy Sciences. HeLa cells were plated onto coverslips coated with laminin or 0.4 μm 12-mm Transwell-COL inserts (Corning) and allowed to adhere overnight. Transwells were washed once with warm PBS and fixed overnight at 4°C with 3% glutaraldehyde in SEM buffer (0.1 M sucrose and 0.1 M Na-phosphate, pH 7.4). Samples were washed with SEM buffer, postfixed with 1% OsO_4_ in SEM buffer on ice for 1 hr, and washed with SEM buffer. Samples were then dehydrated in a graded ethanol series, dried with hexamethyldisilazane, mounted on aluminum stubs, and coated with gold/platinum in a sputter coater. Coverslips were washed with 0.1 M HEPES (pH 7.3) and then fixed with 4% paraformaldehyde, 2.5% glutaraldehyde, and 2 mM CaCL_2_ in 0.1M HEPES for 1 hour at RT. Samples were washed five times with 0.1 M HEPES and incubated with 1% tannic acid in HEPES buffer for 1 hour. Samples were then washed five times with ddH_2_O and incubated with 1% OsO_4_ in ddH_2_O. Samples were washed five times with ddH_2_O, incubated with 1% uranyl acetate in ddH_2_O for 30 minutes, and washed with ddH_2_O. Samples were then dehydrated in a graded ethanol series and dried using critical point drying. Coverslips were mounted on aluminum stubs and coated with gold/palladium using a sputter coater. Imaging was performed using a Quanta 250 Environmental scanning electron microscope operated in high vacuum mode with an accelerating voltage of 5-10 kV. Images were contrast enhanced and cropped using ImageJ software (NIH).

### Image analysis and statistical testing

All digital image analysis was performed using ImageJ or Nikon Elements. Tip enrichment ratios were calculated similar to what we described previously (47). Briefly, confocal images of transfected HeLa cells were used to measure the raw 16-bit fluorescence intensity of a given puncta at the tip of a filopodium; this value was then divided by the cytoplasmic intensity measured immediately at the base of that protrusion. Measurements were typically performed on a minimum of 10 protrusions per cell, from multiple cells cotransfected in at least two independent experiments. For all figures, error bars indicate SD and the number of individual filopodia scored is reported in the legends. All graphs were generated and statistical analyses performed in Prism v.6 or 7 (GraphPad). Unpaired t-test were used to determine levels of statistical significance.

## ACKNOWLEDGEMENTS

We thank all members of the Tyska laboratory, Vanderbilt Microtubule and Motors Club, Vanderbilt Epithelial Biology Center, and Vanderbilt Program in Developmental Biology for advice and support. Super-resolution and electron microcopy were performed through the use of the Vanderbilt Cell Imaging Shared Resource. This work was supported by the Vanderbilt Training Program in Stem Cell and Regenerative Developmental Biology (M.L.W.), American Heart Association Predoctoral Fellowship (M.L.W.), NRSA Predoctoral Fellowship F31DK108528 (M.L.W.), American Heart Association Postdoctoral Fellowship (S.W.C.), and National Institutes of Health Grants R01-DK111949 and R01-DK095811 (M.J.T.). The authors declare no competing financial interests.

## AUTHOR CONTRIBUTIONS

M.L.W. and M.J.T. designed experiments, analyzed data, and wrote the manuscript. M.L.W. performed experiments and MJ.T. supervised the study. S.W.C. assisted with data collection and reagents, respectively. All authors contributed to editing the manuscript.

---

**Supplemental Fig. 1.**
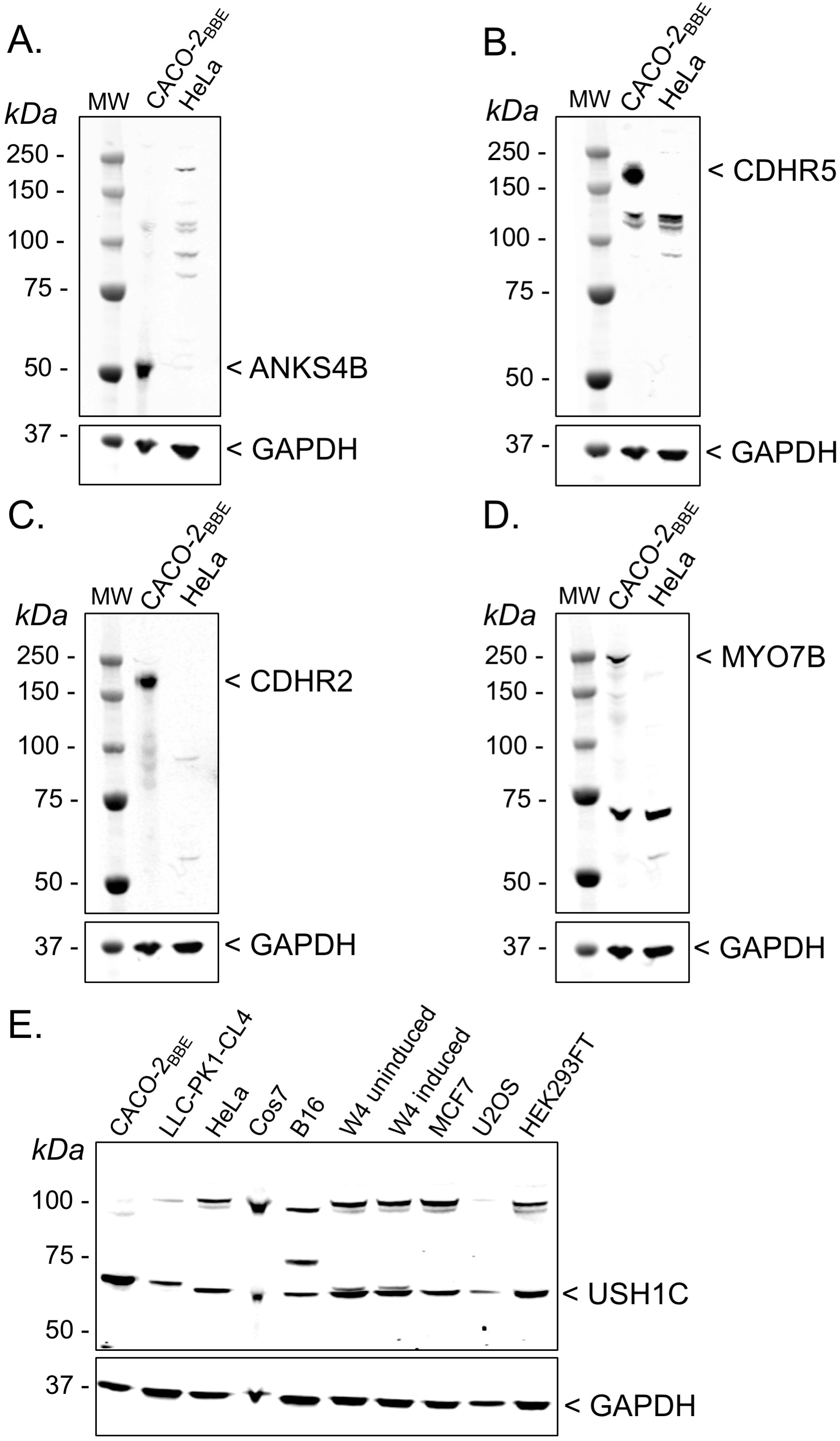

## REFERENCES

1. Pelaseyed, T., and Bretscher, A. (2018) Regulation of actin-based apical structures on epithelial cells. Journal of cell science 131

2. Crawley, S. W., Mooseker, M. S., and Tyska, M. J. (2014) Shaping the intestinal brush border. The Journal of cell biology 207, 441–451

3. Delacour, D., Salomon, J., Robine, S., and Louvard, D. (2016) Plasticity of the brush border - the yin and yang of intestinal homeostasis. Nat Rev Gastroenterol Hepatol 13, 161–174

4. Helander, H. F., and Fandriks, L. (2014) Surface area of the digestive tract - revisited. Scandinavian journal of gastroenterology

5. Mooseker, M. S., and Tilney, L. G. (1975) Organization of an actin filament-membrane complex. Filament polarity and membrane attachment in the microvilli of intestinal epithelial cells. The Journal of cell biology 67, 725–743

6. Tyska, M. J., Mackey, A. T., Huang, J. D., Copeland, N. G., Jenkins, N. A., and Mooseker, M. S. (2005) Myosin-1a is critical for normal brush border structure and composition. Molecular biology of the cell 16, 2443–2457

7. Pinette, J. A., Mao, S., Millis, B. A., Krystofiak, E. S., Faust, J. J., and Tyska, M. J. (2019) Brush border protocadherin CDHR2 promotes the elongation and maximized packing of microvilli in vivo. Molecular biology of the cell 30, 108–118

8. Li, J., He, Y., Weck, M. L., Lu, Q., Tyska, M. J., and Zhang, M. (2017) Structure of Myo7b/USH1C complex suggests a general PDZ domain binding mode by MyTH4-FERM myosins. Proceedings of the National Academy of Sciences of the United States of America 114, E3776–E3785

9. Weck, M. L., Crawley, S. W., Stone, C. R., and Tyska, M. J. (2016) Myosin-7b Promotes Distal Tip Localization of the Intermicrovillar Adhesion Complex. Current biology: CB 26, 2717–2728

10. Crawley, S. W., Weck, M. L., Grega-Larson, N. E., Shifrin, D. A., Jr., and Tyska, M. J. (2016) ANKS4B Is Essential for Intermicrovillar Adhesion Complex Formation. Dev Cell 36, 190–200

11. Crawley, S. W., Shifrin, D. A., Jr., Grega-Larson, N. E., McConnell, R. E., Benesh, A. E., Mao, S., Zheng, Y., Zheng, Q. Y., Nam, K. T., Millis, B. A., Kachar, B., and Tyska, M. J. (2014) Intestinal brush border assembly driven by protocadherin-based intermicrovillar adhesion. Cell 157, 433–446

12. Yu, I. M., Planelles-Herrero, V. J., Sourigues, Y., Moussaoui, D., Sirkia, H., Kikuti, C., Stroebel, D., Titus, M. A., and Houdusse, A. (2017) Myosin 7 and its adaptors link cadherins to actin. Nat Commun 8, 15864

13. Li, J., He, Y., Lu, Q., and Zhang, M. (2016) Mechanistic Basis of Organization of the Harmonin/USH1C-Mediated Brush Border Microvilli Tip-Link Complex. Dev Cell 36, 179–189

14. Choi, M. S., Graves, M. J., Matoo, S., Storad, Z. A., El Sheikh Idris, R. A., Weck, M. L., Smith, Z. B., Tyska, M. J., and Crawley, S. W. (2020) The small EF-hand protein CALML4 functions as a critical myosin light chain within the intermicrovillar adhesion complex. J Biol Chem 295, 9281–9296

15. Siemens, J., Lillo, C., Dumont, R. A., Reynolds, A., Williams, D. S., Gillespie, P. G., and Muller, U. (2004) Cadherin 23 is a component of the tip link in hair-cell stereocilia. Nature 428, 950–955

16. Ahmed, Z. M., Goodyear, R., Riazuddin, S., Lagziel, A., Legan, P. K., Behra, M., Burgess, S. M., Lilley, K. S., Wilcox, E. R., Riazuddin, S., Griffith, A. J., Frolenkov, G. I., Belyantseva, I. A., Richardson, G. P., and Friedman, T. B. (2006) The tip-link antigen, a protein associated with the transduction complex of sensory hair cells, is protocadherin-15. J Neurosci 26, 7022–7034

17. Kazmierczak, P., Sakaguchi, H., Tokita, J., Wilson-Kubalek, E. M., Milligan, R. A., Muller, U., and Kachar, B. (2007) Cadherin 23 and protocadherin 15 interact to form tip-link filaments in sensory hair cells. Nature 449, 87–91

18. Siemens, J., Kazmierczak, P., Reynolds, A., Sticker, M., Littlewood-Evans, A., and Muller, U. (2002) The Usher syndrome proteins cadherin 23 and harmonin form a complex by means of PDZ-domain interactions. Proc Natl Acad Sci USA 99, 14946–14951

19. Boeda, B., El-Amraoui, A., Bahloul, A., Goodyear, R., Daviet, L., Blanchard, S., Perfettini, I., Fath, K. R., Shorte, S., Reiners, J., Houdusse, A., Legrain, P., Wolfrum, U., Richardson, G., and Petit, C. (2002) Myosin VIIa, harmonin and cadherin 23, three Usher I gene products that cooperate to shape the sensory hair cell bundle. EMBO J 21, 6689–6699

20. Weil, D., El-Amraoui, A., Masmoudi, S., Mustapha, M., Kikkawa, Y., Laine, S., Delmaghani, S., Adato, A., Nadifi, S., Zina, Z. B., Hamel, C., Gal, A., Ayadi, H., Yonekawa, H., and Petit, C. (2003) Usher syndrome type I G (USH1G) is caused by mutations in the gene encoding SANS, a protein that associates with the USH1C protein, harmonin. Hum Mol Genet 12, 463–471

21. Adato, A., Michel, V., Kikkawa, Y., Reiners, J., Alagramam, K. N., Weil, D., Yonekawa, H., Wolfrum, U., El-Amraoui, A., and Petit, C. (2005) Interactions in the network of Usher syndrome type 1 proteins. Hum Mol Genet 14, 347–356

22. Senften, M., Schwander, M., Kazmierczak, P., Lillo, C., Shin, J. B., Hasson, T., Geleoc, G. S., Gillespie, P. G., Williams, D., Holt, J. R., and Muller, U. (2006) Physical and functional interaction between protocadherin 15 and myosin VIIa in mechanosensory hair cells. J Neurosci 26, 2060–2071

23. Pan, L., Yan, J., Wu, L., and Zhang, M. (2009) Assembling stable hair cell tip link complex via multidentate interactions between harmonin and cadherin 23. Proc Natl Acad Sci USA 106, 5575–5580

24. Bahloul, A., Michel, V., Hardelin, J. P., Nouaille, S., Hoos, S., Houdusse, A., England, P., and Petit, C. (2010) Cadherin-23, myosin VIIa and harmonin, encoded by Usher syndrome type I genes, form a ternary complex and interact with membrane phospholipids. Hum Mol Genet 19, 3557–3565

25. Yan, J., Pan, L., Chen, X., Wu, L., and Zhang, M. (2010) The structure of the harmonin/sans complex reveals an unexpected interaction mode of the two Usher syndrome proteins. Proc Natl Acad Sci U S A 107, 4040–4045

26. Grati, M., and Kachar, B. (2011) Myosin VIIa and sans localization at stereocilia upper tip-link density implicates these Usher syndrome proteins in mechanotransduction. Proceedings of the National Academy of Sciences of the United States of America 108, 11476–11481

27. Lentz, J., and Keats, B. J. B. (1993) Usher Syndrome Type I. in GeneReviews(R) (Pagon, R. A., Adam, M. P., Ardinger, H. H., Wallace, S. E., Amemiya, A., Bean, L. J. H., Bird, T. D., Fong, C. T., Mefford, H. C., Smith, R. J. H., and Stephens, K. eds.), Seattle (WA). pp

28. Schwander, M., Kachar, B., and Muller, U. (2010) Review series: The cell biology of hearing. The Journal of cell biology 190, 9–20

29. Bitner-Glindzicz, M., Lindley, K. J., Rutland, P., Blaydon, D., Smith, V. V., Milla, P. J., Hussain, K., Furth-Lavi, J., Cosgrove, K. E., Shepherd, R. M., Barnes, P. D., O’Brien, R. E., Farndon, P. A., Sowden, J., Liu, X. Z., Scanlan, M. J., Malcolm, S., Dunne, M. J., Aynsley-Green, A., and Glaser, B. (2000) A recessive contiguous gene deletion causing infantile hyperinsulinism, enteropathy and deafness identifies the Usher type 1C gene. Nature genetics 26, 56–60

30. Verpy, E., Leibovici, M., Zwaenepoel, I., Liu, X. Z., Gal, A., Salem, N., Mansour, A., Blanchard, S., Kobayashi, I., Keats, B. J., Slim, R., and Petit, C. (2000) A defect in harmonin, a PDZ domaincontaining protein expressed in the inner ear sensory hair cells, underlies Usher syndrome type 1C. Nature genetics 26, 51–55

31. Pena, J. F., Alie, A., Richter, D. J., Wang, L., Funayama, N., and Nichols, S. A. (2016) Conserved expression of vertebrate microvillar gene homologs in choanocytes of freshwater sponges. Evodevo 7, 13

32. He, Y., Li, J., and Zhang, M. (2019) Myosin VII, USH1C, and ANKS4B or USH1G Together Form Condensed Molecular Assembly via Liquid-Liquid Phase Separation. Cell Rep 29, 974–986 e974

33. Henn, A., and De La Cruz, E. M. (2005) Vertebrate myosin VIIb is a high duty ratio motor adapted for generating and maintaining tension. The Journal of biological chemistry 280, 39665–39676

34. Yang, Y., Kovacs, M., Xu, Q., Anderson, J. B., and Sellers, J. R. (2005) Myosin VIIB from Drosophila is a high duty ratio motor. The Journal of biological chemistry 280, 32061–32068

35. Bird, J. E., Barzik, M., Drummond, M. C., Sutton, D. C., Goodman, S. M., Morozko, E. L., Cole, S. M., Boukhvalova, A. K., Skidmore, J., Syam, D., Wilson, E. A., Fitzgerald, T., Rehman, A. U., Martin, D. M., Boger, E. T., Belyantseva, I. A., and Friedman, T. B. (2017) Harnessing molecular motors for nanoscale pulldown in live cells. Molecular biology of the cell 28, 463–475

36. Bohil, A. B., Robertson, B. W., and Cheney, R. E. (2006) Myosin-X is a molecular motor that functions in filopodia formation. Proceedings of the National Academy of Sciences of the United States of America 103, 12411–12416

37. Zhang, H., Berg, J. S., Li, Z., Wang, Y., Lang, P., Sousa, A. D., Bhaskar, A., Cheney, R. E., and Stromblad, S. (2004) Myosin-X provides a motor-based link between integrins and the cytoskeleton. Nature cell biology 6, 523–531

38. Berg, J. S., and Cheney, R. E. (2002) Myosin-X is an unconventional myosin that undergoes intrafilopodial motility. Nature cell biology 4, 246–250

39. Kerber, M. L., Jacobs, D. T., Campagnola, L., Dunn, B. D., Yin, T., Sousa, A. D., Quintero, O. A., and Cheney, R. E. (2009) A novel form of motility in filopodia revealed by imaging myosin-X at the single-molecule level. Current biology: CB 19, 967–973

40. Belyantseva, I. A., Boger, E. T., Naz, S., Frolenkov, G. I., Sellers, J. R., Ahmed, Z. M., Griffith, A. J., and Friedman, T. B. (2005) Myosin-XVa is required for tip localization of whirlin and differential elongation of hair-cell stereocilia. Nature cell biology 7, 148–156

41. Marshall, W. F. (2015) How Cells Measure Length on Subcellular Scales. Trends Cell Biol 25, 760–768

42. Burke, T. A., Christensen, J. R., Barone, E., Suarez, C., Sirotkin, V., and Kovar, D. R. (2014) Homeostatic actin cytoskeleton networks are regulated by assembly factor competition for monomers. Current biology: CB 24, 579–585

43. Rotty, J. D., Wu, C., Haynes, E. M., Suarez, C., Winkelman, J. D., Johnson, H. E., Haugh, J. M., Kovar, D. R., and Bear, J. E. (2015) Profilin-1 serves as a gatekeeper for actin assembly by Arp2/3-dependent and -independent pathways. Developmental cell 32, 54–67

44. Suarez, C., Carroll, R. T., Burke, T. A., Christensen, J. R., Bestul, A. J., Sees, J. A., James, M. L., Sirotkin, V., and Kovar, D. R. (2015) Profilin regulates F-actin network homeostasis by favoring formin over Arp2/3 complex. Developmental cell 32, 43–53

45. Suarez, C., and Kovar, D. R. (2016) Internetwork competition for monomers governs actin cytoskeleton organization. Nature reviews. Molecular cell biology 17, 799–810

46. Faust, J. J., Millis, B. A., and Tyska, M. J. (2019) Profilin-Mediated Actin Allocation Regulates the Growth of Epithelial Microvilli. Curr Biol 29, 3457–3465 e3453

47. Mazerik, J. N., and Tyska, M. J. (2012) Myosin-1A targets to microvilli using multiple membrane binding motifs in the tail homology 1 (TH1) domain. J Biol Chem 287, 13104–13115

